## Supplementary Files for "Transgenic expression of *cif* genes from *Wolbachia* strain *w*AlbB recapitulates cytoplasmic incompatibility in *Aedes aegypti*"

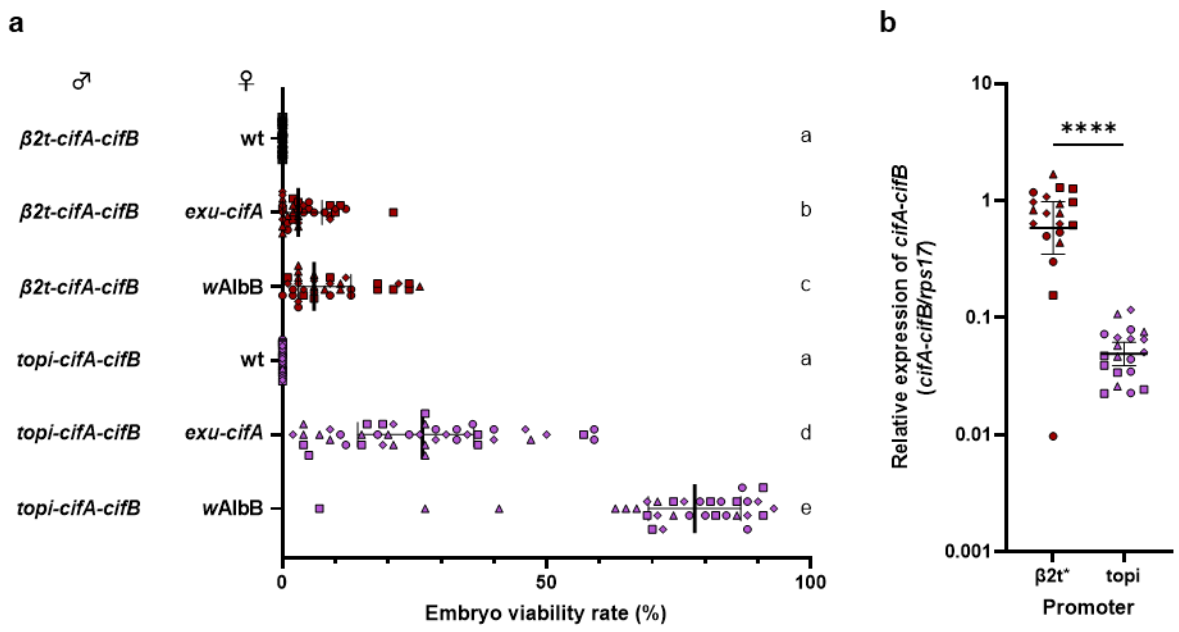


**Supplementary Fig. 1 | Results of incompatibility crosses were consistent between different genomic insertion lines. a,** Lowering the dosage of *cifA-cifB* increases the rescue capability of either transgenic (*exu*-cifA) or *w*AlbB-carrying females. **b,** The relative expression of *cifA-cifB* was higher when controlled by the *β2t* as opposed to the *topi* promoter (unpaired t-test $p<0.0001$, mean and s.d. are shown). In both **a,** and **b,** circle, square, triangle and diamond symbols represent biological replicates from *β2t-cifA-cifB* and *topi-cifA-cifB* insertion sites 1,2,3, and 4 respectively.

**Supplementary Video 1. Timelapse of wild-type and *β2t-cifB* male reproductive tissues.** Mature spermatozoa were released (black arrow) from squished reproductive tissues dissected from wild-type (WT) males but not *β2t-cifB* males, tissues were imaged at 10x magnification.
